# Temporal Dynamics of Cyanobacterial Bloom Community Composition and Toxin Production from Urban Lakes

**DOI:** 10.1101/2024.02.07.579333

**Authors:** Julie A. Maurer, Andrew M. Kim, Nana Oblie, Sierra Hefferan, Hannuo Xie, Angela Slitt, Bethany D. Jenkins, Matthew J. Bertin

## Abstract

With a long evolutionary history and a need to adapt to a changing environment, cyanobacteria in freshwater systems use specialized metabolites for communication, defense, and physiological processes. However, the role that these metabolites play in differentiating species, maintaining microbial communities, and generating niche persistence and expansion is poorly understood. Furthermore, many cyanobacterial specialized metabolites and toxins present significant human health concerns due to their liver toxicity and their potential impact to drinking water. Gaps in knowledge exist with respect to changes in species diversity and toxin production during a cyanobacterial bloom (cyanoHAB) event; addressing these gaps will improve understanding of impacts to public and ecological health. In the current project, we utilized a multi-omics strategy (DNA metabarcoding and metabolomics) to determine the cyanobacterial community composition, toxin profile, and the specialized metabolite pool at three freshwater lakes in Providence, RI during summer-fall cyanoHABs. Species diversity decreased at all study sites over the course of the bloom event, and toxin production reached a maximum at the midpoint of the event. Additionally, LC-MS/MS-based molecular networking identified new toxin congeners. This work provokes intriguing questions with respect to the use of allelopathy by organisms in these systems and the presence of emerging toxic compounds that can impact public health.

**SYNOPSIS:** This study reports on cyanobacterial community succession and toxin dynamics during cyanobacterial bloom events. Results show relationships and temporal dynamics that are relevant to public health.

## INTRODUCTION

Cyanobacterial blooms (cyanoHABs) are complex microbial systems subject to alterations in abiotic and biotic parameters. These blooms produce a suite of alkaloidal neurotoxins and peptidic hepatotoxins, which can cause non-alcoholic fatty liver disease and acute liver failure.^1,2^ Additionally, these events pose significant risks to freshwater drinking water resources, perhaps most vividly exemplified by the Toledo Water Crisis of 2014, in which levels of the hepatotoxic microcystin-LR (MC-LR) in drinking water led to a “do not drink; do not use” warning.^3^ The majority of research and environmental monitoring has been devoted to microcystin-LR, and concentration limits are set by the US EPA (1.6 μg/L = Do Not Drink; 20 μg/L = Do Not Use). The World Health Organization only sets exposure guidelines for this one microcystin variant (MC-LR) and three alkaloidal toxins (anatoxin-a, saxitoxin, and cylindrospermopsin).^4^ However, 2,425 cyanobacterial metabolites have been described to date, including 318 microcystins along with other cyanopeptides such as the anabaenopeptins, cyanopeptolins, and microginins.^5^

Understanding the full suite of toxins and the temporal resolution of their production is key to determining the full human health impact of these events and the impact of these metabolites on shaping bacterial communities. CyanoHABs represent a complex mixture of species and toxins. Many species in these events have different toxin biosynthesis capabilities, and the toxins produced vary in potency and biological mechanism of action.^6^ Water monitoring and treatment can only account for toxins that are currently known, and many testing methods are not able to distinguish between microcystin congeners of varying potency. These additional toxins (AKA emerging toxins, ETs) would not be monitored in raw water, treated water, or finished water. Our group and collaborators have previously used a large-scale harvesting platform to access unprecedented amounts of cyanobacteria biomass, leading to chemical extraction, fractionation, metabolite profiling, and isolation procedures. This research enabled the identification of new exquisitely potent cytotoxic metabolites (the cyanobufalins) and new potently cytotoxic microcystins (MCs).^7,8^

The cyanobacterial communities in these systems change due to nitrogen source (nitrogen fixers vs. non nitrogen fixers),^9^ but they also exhibit successional patterns based on season and temperature (summer blooms vs. fall blooms).^10^ For instance, *Microcystis* species have been documented to dominate during summer blooms, while other taxa such as *Aphanizomenon* are present during cooler weather.^11^ The specter of a changing climate bodes for increases in cyanoHABs in freshwater bodies,^12^ with worsening summer blooms and a potential for toxic blooms later in the fall. Increasing temperatures and nutrient regimes will impact these these existing community dynamics and may alter the composition of the cyanotoxin pool during bloom events.^13^

The goal of the current work was to determine cyanobacterial community composition and toxin production alterations over time during cyanoHAB events at three urban lakes using DNA metabarcoding and untargeted metabolomics approaches. The data showed that communities shifted over time with decreasing species diversity throughout the event culminating in domination by *Aphanizomenon*. Additionally, cyanotoxins did not show constitutive production but appeared in a punctuated fashion correlating strongly with cyanobacterial biomass. As the bloom progressed, metabolite profiles shifted suggested the production of a unique metabolite pool as species succession occurred. This work identified putative new cyanotoxins and set baseline time series data on community composition that can be augmented by future field studies. Furthermore, the data presented drive new hypotheses with respect to the effects of climate change on community composition and toxin production.

## METHODS AND MATERIALS

### Sample Sites and Sample Collection

Surface water samples were acquired weekly over the course of a cyanoHAB event at Roger Williams Park in Providence, Rhode Island (Figure S1). The cyanoHAB event was observed in three lakes at the park (Polo Lake, Pleasure Lake, and Cunliff Lake) from September 1, 2022 to December 7, 2022. One surface water sample was taken from Blackstone Pond in Cranston, RI on 9/8/2022, but weekly sampling did not continue at this site. The water samples were transported to the laboratory and chlorophyll a values were measured using UV absorbance following filtration of 50 mL of surface water over a GF/F filter and extraction with methanol.^14^ Additional samples (30 mL) were filtered over a 47 mm 0.2 μm Sterlitech polyester track-etch (PETE) filter and stored at -80 °C for subsequent DNA extraction. A third portion of surface water (50 mL) was filtered over a 47 mm 3 μm PETE filter to capture cyanobacterial cells for toxin analysis. These filters were stored at -20 °C prior to analysis. In all, a total of 43, 46, and 44 samples were analyzed for chlorophyll a, cyanobacterial community, and toxin composition, respectively.

### DNA Isolation and Metabarcoding

DNA was extracted from biomass collected on the 47 mm 0.2 μm Sterlitech PETE filters using a modified version of the Qiagen Plant MiniKit with 4 μL RNase-A added, a 1-minute bead-beating step, and a total elution volume of 30 μL Buffer AE. For each sample, DNA was quantified using a Nanodrop 2000 prior to amplification. Gene amplifications were completed using cyanobacteria-specific primers^15^ with MiSeq adapters to target a region of the 16S rRNA gene between target sites at 359 and 701 bps (Primers - CYA359 forward primer: 5’ GAA TTT TCC GCA ATG GG 3’; CYA781Ra reverse primer: 5’ GAC TAC TGG GGT ATC TAA TCC CAT T 3’; CYA781Rb reverse primer: 5’ GAC TAC AGG GGT ATC TAA TCC CTT T 3’). Amplicons were then sequenced using Illumina MiSeq Next-Generation Sequencing platform using V3 chemistry and 2 × 250 bp paired-end sequencing at the University of Rhode Island Molecular Informatics Core. Sequences were returned as de-multiplexed reads. Raw reads were quality controlled before and after trimming with FastQC (v0.11.7)^16^ and MultiQC (v1.14).^17^ MiSeq adapters and primers were trimmed using CutAdapt (v4.4).^18^ DADA2 (v1.26.0)^19^ was used in the coding platform R to estimate sequencing error and to determine amplicon sequence variants (ASVs). Taxonomy was assigned using the SILVA reference database (v138.1)^20^ and NCBI BLAST and ASVs determined to be chloroplast DNA, mitochondrial DNA, and unassigned sequences were removed. The processed dataset of 107 ASVs across 46 samples was then imported into Phyloseq (v. 1.44.0),^21^ an R package for visualization and analysis of molecular sequence data. Phyloseq was used to perform zero-replacement^22^ and center-log ratio (CLR) transformations on raw read abundances in accordance with current best practices for analyzing compositional data (e.g., bacterial sequences).^23^ Aitchison distances were then calculated prior to Non-metric MultiDimensional Scaling (NMDS) ordination and Analysis of Similarities (ANOSIM). Patterns in cyanobacterial species beta diversity across sample sites were examined with NMDS ordination techniques and quantitatively tested for group dispersion (permutest, R) and analysis of variance among groups (ANOSIM, PERMANOVA, R). Alpha diversity trends over time were calculated with chao1 and shannon index^24^ and trends in alpha diversity over time were quantitatively assessed with the Mann-Kendall test, a statistical test for time series trends.^25^ To qualitatively examine successional patterns of cyanobacterial species over time, raw read abundances were also transformed to relative abundances and plotted in phyloseq. All samples were used for the NMDS plots for every location (Cunliff, Polo, and Pleasure) to examine community composition by site. The raw sequence reads have been deposited to the Sequence Read Archive at Genbank (BioProject #: PRJNA1065459), which will be available upon publication.

### Toxin analysis

The 47 mm 3 μm filters were thawed at room temperature and repeatedly extracted in 70%/25%/5% CH_3_OH, H_2_O, n-butanol until no color was observed in the extract. Extracts for each sample were pooled and then concentrated *in vacuo*, reconstituted in CH_3_OH and subjected to a sample preparation step whereby each extract was passed over a 100 mg C18 SPE cartridge eluting with 1 mL of CH_3_OH directly into an LC-MS vial. Each sample was analyzed via LC-MS/MS to determine cyanotoxin composition. Raw mass spectrometry data were collected using a Dionex Ultimate 3000 HPLC system coupled to a Thermo Scientific LTQ XL mass spectrometer. A 2.6 μm Kinetex C18 column (150 × 4.6 mm) was used for separations with a flow rate set at 0.4 mL/min. A gradient method consisting of H_2_O with 0.1% formic acid and CH_3_CN with 0.1% formic acid was used. The gradient method was as follows: 15% CH_3_CN was held for 5 min, followed by a 15 min gradient to 100% CH_3_CN, which was held for 10 min followed by a return to initial conditions from 31-38 min. For the data dependent MS/MS component, the CID isolation width was 1.0 and the collision energy was 35.0 eV.

### LC-MS/MS-based Molecular Networking

Once acquisitions were complete, .raw data files were exported and transformed into .mzXML format using MSConvert.^26^ Files were uploaded into the Global Natural Products Social Molecular Networking platform (GNPS).^27^ Network parameters included a parent mass tolerance and MS/MS fragment mass tolerance of 2.0 Da and 0.5 Da, respectively for low resolution data. The network’s cosine score for edge connection was set to 0.6 with at least 3 matched peaks required. The spectra were searched against the GNPS library of spectra, where matched nodes needed to have at least 6 matched peaks and meet the minimum cosine score of 0.7. Library hits were annotated for each sample and verified by MS/MS fragmentation comparisons. Cyanotoxins were annotated using the GNPS platform either by specific library hits or by matches to MS/MS spectra from literature sources. Precursor *m/z* values of identified cyanotoxins were extracted using the GNPS Dashboard function and abundance values were annotated for each toxin.^28^ To garner total metabolite counts for each sample, each file was uploaded to MSDIAL,^29^ and peak lists were generated for each MS acquisition to give a list of metabolites. Correlations were performed by computing Pearson correlation coefficients in Prism (v. 9.5.1). All raw mass spectrometry files and .mzXML files are available in the MassIVE repository (accession#: MSV000093676).

### Principal Component Analysis

The peak lists for all samples were aligned, and peak list tables including retention time, mass to charge ratio, and peak intensity were exported as .csv files. The peak lists were uploaded to MetaboAnalyst (v. 2.0),^30^ and the default parameters for PCA analysis were used. Data were then filtered by mean intensity value and the data were Pareto scaled (mean-centered and divided by the square root of the standard deviation of each variable). Finally, PCA plots were generated to examine similarities in metabolite composition at all sites and with respect to two time periods (early: 9/1/2022-10/19/2022; late: 10/26/2022-12/7/2022).

## RESULTS AND DISCUSSION

### Fall CyanoHABs in Urban Lakes

CyanoHABs were observed at the three study sites beginning on September 1, 2022, and cyanobacterial cell density increased over time in these water bodies as observed by photography, analysis of water samples by microscopy, and by examination of chlorophyll a values (Figures S2-S5). Biomass maxima were reached on November 2, 2022, at Pleasure and Cunliff Lakes, while the biomass maximum was reached on October 19, 2022, at Polo Lake (Figure S5). We ended sampling on December 14, 2022, after seeing no cyanobacterial cells in surface water samples from two sites (Pleasure and Cunliff Lakes). Surface samples were processed in the laboratory for chlorophyll a analysis, DNA metabarcoding, and toxin analysis. *Microcystis aeruginosa* NIES-933 (UTEX LB 2385) was purchased from the UTEX culture collection (a known MC-LR producer) and served as a positive control for both molecular sequencing and toxin analysis. Following microscopy analysis of environmental samples, community shifts were noted from early to late bloom periods with the identification of *Aphanizomenon* colonies in late November and December samples (Figure S4). These observations were further supported by subsequent molecular analysis (see below).

### Cyanobacterial Species Distribution across Space and Time

Qualitative examination of cyanobacterial relative abundances at the genus level showed a distinct successional pattern over the study period at all three sites. Diverse filamentous species such as *Cuspidothrix* sp., *Dolichospermum* sp., and *Planktothrix* sp. were observed during the first two weeks of sampling with *Microcystis* and *Woronichinia* sp. (family Microcystaceae) co-dominating the remainder of the earlier and middle bloom periods (9/14/2022 – 11/2022). *Aphanizomenon* sp. (family Nostocaceae) showed the highest relative abundance during the later bloom, with complete community dominance by 11/30/2022 (Figure 1). Cunliff Lake showed lower relative levels of *Woronichinia* compared to the other two lakes and the presence and relative abundance of *Aphanizomenon* sp. occurred earlier (Figure 1). NMDS ordination showed enough separation between the composition of the cyanobacterial community between September and October - December (stress = 0.132, dimensions = 2) to delineate two succession “stages” in the bloom, which we classified as “early” and “late” (Figure S6). This separation was supported by tests of variance between the grouping of succession and genus (ANOSIM, p < 0.001, R = 0.586).

**Figure 1.**
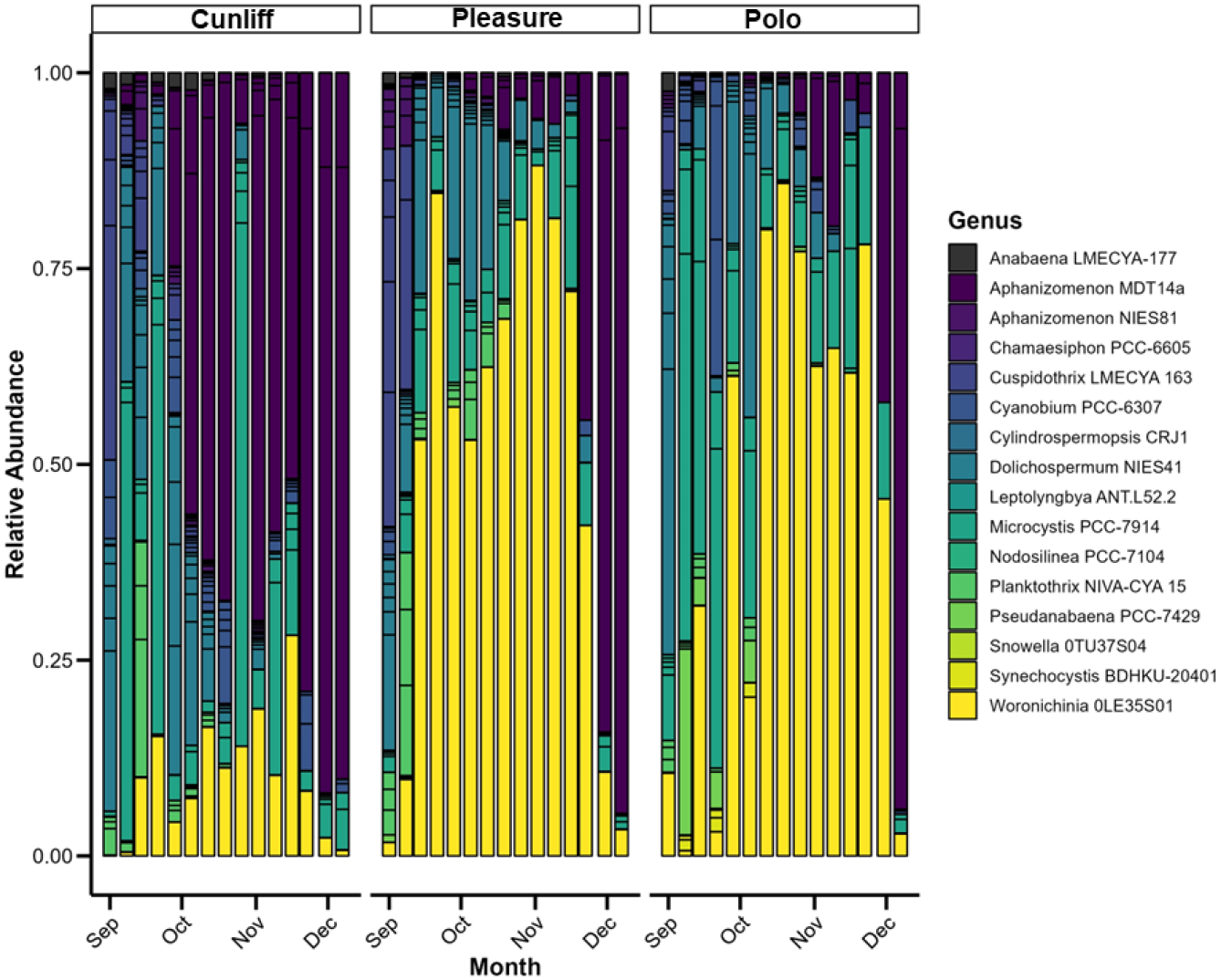
CyanoHAB community composition and successional patterns. Relative abundances of cyanobacterial taxa at Cunliff, Pleasure, and Polo Lakes from 9/1/2022 to 12/7/2022.

However, permutation tests showed significant dispersion within the succession grouping (p < 0.05), and the succession groups identified still contained some overlap and a high degree of scatter. Sample sites differed in taxonomic composition (ANOSIM, R = 0.14, p < 0.01), with this difference driven by Cunliff Lake, though again permutation tests revealed a significant amount of dispersion (p < 0.01) (Figure S7). Examining cyanobacterial genera by site, Cunliff Lake was the most unique and was dominated more by *Microcystis* than *Woronichinia* in the early bloom period and then dominated by *Aphanizomenon* earlier in the time series compared to Polo and Pleasure Lakes (Figure 1). The positive control (UTEX LB 2385, *M. aeruginosa* NIES-933) ASVs were identified as 100% *Microcystis* following our bioinformatics analysis, and the sequence match was 100% to *M. aeruginosa* NIES 933 (Figure S8A). Additionally, MC-LR was detected in extracts of this strain with an MS/MS match to that of the MC-LR in the GNPS library (Figure S8B).

The cyanobacterial community was more diverse at the onset of the bloom and diversity decreased as the bloom progressed by both chao1 (richness) and shannon (richness and evenness) metrics across all sample sites (Figure 2). Mann-Kendall test results showed that these decreases over time were significant across all metrics (chao: tau =-0.61, p < 0.001; shannon: tau = -0.19, p < 0.001). The use of DNA barcoding for community analysis instead of cell counts likely increased the species resolution over the course of the bloom and allowed for a clear picture of decreasing species diversity as the cyanoHABs progressed.

**Figure 2.**
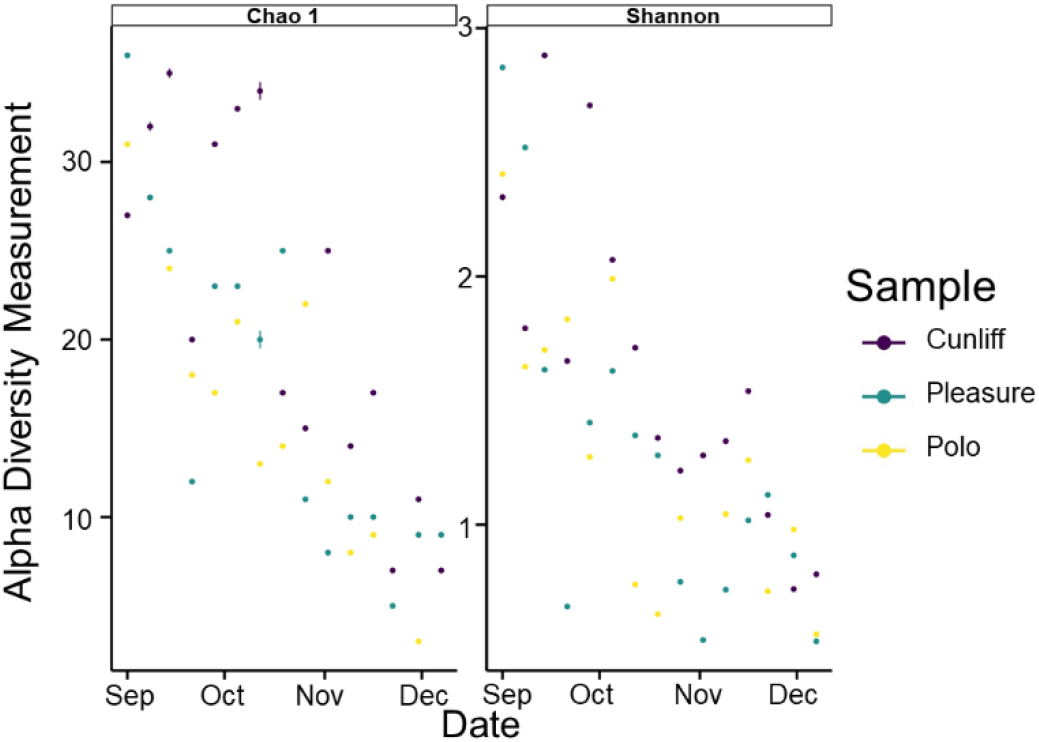
Species diversity decreased over the course of cyanoHABs. Alpha diversity measurements completed over the time period 9/1/2022 to 12/7/2022 determined by Chao and Shannon metrics.

The molecular community composition data are consistent with past work done using cell counts and cultivation studies. Multiple studies have shown changing cyanobacterial successional patterns either from *Aphanizomenon* sp. to *Microcystis* or vice versa likely driven by temperature and this pattern can oscillate seasonally.^9,10,31,32^ Culture studies have shown that *Microcystis* sp. has a higher maximum growth rate at warmer temperatures than *Aphanizomenon* sp.^33^ In addition to temperature, nitrogen source and N:P ratio play significant roles in shaping cyanobacterial community succession (review in Tanvir et al. 2021).^10^ In this study, nutrient data were obtained for Polo Lake, but not Cunliff and Pleasure. Previous studies have shown spatial-temporal variation,^32,34^ while we showed very clear temporal variation and some spatial variation. This may be due to the proximity of the sites used in the study and some connection of water masses amongst the lakes. The study site lakes were likely subjected to similar nutrient regimes and other abiotic and biotic parameters. Our results suggest that more research should be devoted to the genus *Woronichinia*. This group showed abundance throughout most of the bloom period at two lakes, correlated with the relative abundance of *Microcystis* sp. at Pleasure Lake (Figure S9), and its high abundance in urban lakes has been previously reported.^35^ While this genus does not appear to possess MC biosynthetic genes, clusters for the anabaenopeptins and other cyanopeptides have been annotated.^36^ Some toxicity from this group has been reported against invertebrates,^37^ and further investigation of its ecological role and its potential for allelopathic compound production is warranted.

### Cyanotoxin Production during Blooms

We annotated 14 cyanotoxins from LC-MS/MS data. Microcystin-LR, MC-LA, anabaenopeptin A, and ferintoic acid A were identified using the library search function at GNPS and our putative toxin queries were matched to MS/MS fragmentation patterns for library compounds (Figure S10). Microcystin-LR identified in our environmental samples had an identical retention time, precursor *m/z* and MS/MS fragmentation pattern to that of the MC-LR identified in the UTEX positive control (Figure S10). Additional toxins were identified by examining MS/MS networks (e.g., anabaenopeptin B in same cluster as anabaenopeptin A) and by comparing precursor *m/z* values and MS/MS spectra to previously published work.^38-41^ Toxin concentrations (as determine by AUC values from mass spec data) coincided with cyanobacterial biomass maxima at all three lakes (Figure 3 and Figure S11A) and toxin production was not constitutive. Additionally, metabolite content in general increased with cyanobacterial biomass (Figure S11B).

**Figure 3.**
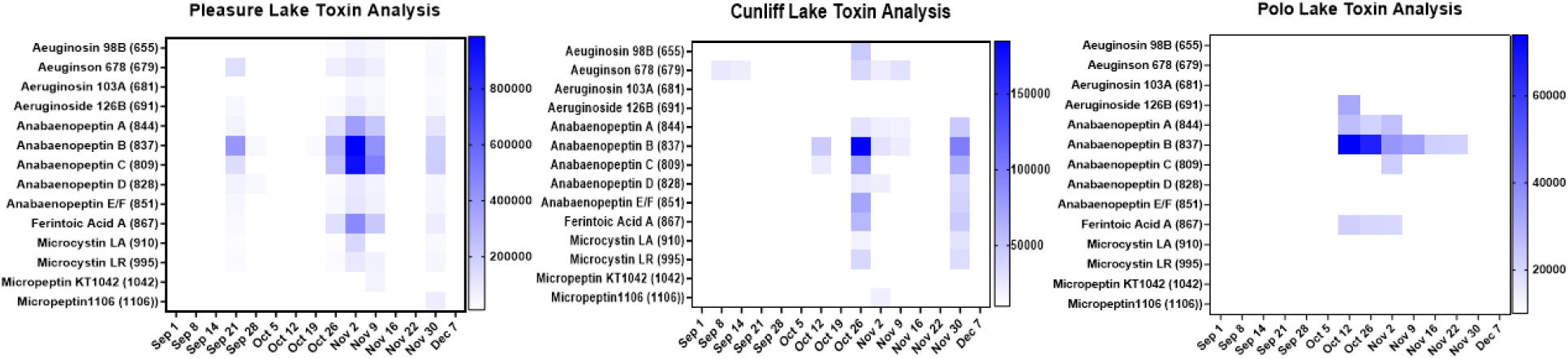
Cyanotoxin annotation and relative concentration at Pleasure, Cunliff, and Polo Lakes during the study period from 9/1/2022 to 12/7/2022. The colors in the heat maps are proportional to the intensity of the extracted ion for each analyte. Precursor masses for each toxin are listed in the parentheses.

Examination of LC-MS/MS networks annotated multiple toxin classes (e.g., microcystins, anabaenopeptins, aeruginosins, and cyanopeptolins) and putative new toxins, but hundreds of metabolites remain to be annotated demonstrating that these events are incredibly chemically complex (Figure 4). The LC-MS/MS networking approach to identify new cyanopeptides and toxins has shown utility,^42^ and this strategy is well suited to define structure-activity relationships amongst toxin classes. Most of the toxins of interest were found at all three lakes (e.g., anabaenopeptins A, B, and ferintoic acid B), but interestingly MC-LR was only detected in samples collected from Pleasure and Cunliff Lakes. Examining metabolite content over the entire sample period and all sites by PCA, we observed very similar composition overall (Figure 5A). However, delineating samples by early and late periods of the bloom, there was a difference in composition driven by the presence and concentration of cyanotoxins (Figure 5B). In terms of connecting specific cyanobacterial taxa to toxin production, there was a high relative abundance of *Microcystis* at Cunliff Lake on 10/26/22 which coincides with the highest abundances of MCs and other cyanotoxins (cf. Figures 1 and 3) and *Microcystis* relative abundance correlated positively with the abundances of aeruginosins, anabaenopeptins, and microcystins at Cunliff Lake (p < 0.01, 0.01, and 0.05, respectively). However, there were high levels of *Microcystis* observed at Polo Lake early in the bloom period with no corresponding MCs detected, which may be explained by a complicated array of potential triggering factors such as nutrient profiles, competition, and grazing, amongst others. The metabolomics data we report were consistent with metatranscriptomics studies that showed *mcy* and other toxin biosynthesis genes were generally expressed during the later part of blooms.^43,44^ However, this picture is complicated by the number of toxin producing taxa present in cyanoHABs and their ecosystem function and population dynamics. Integrating more ‘omics approaches such as metagenomics and metatranscriptomics with the metabolomics data will give greater insight into the ecophysiological roles of cyanotoxins in shaping cyanobacterial and bacterial communities.

**Figure 4.**
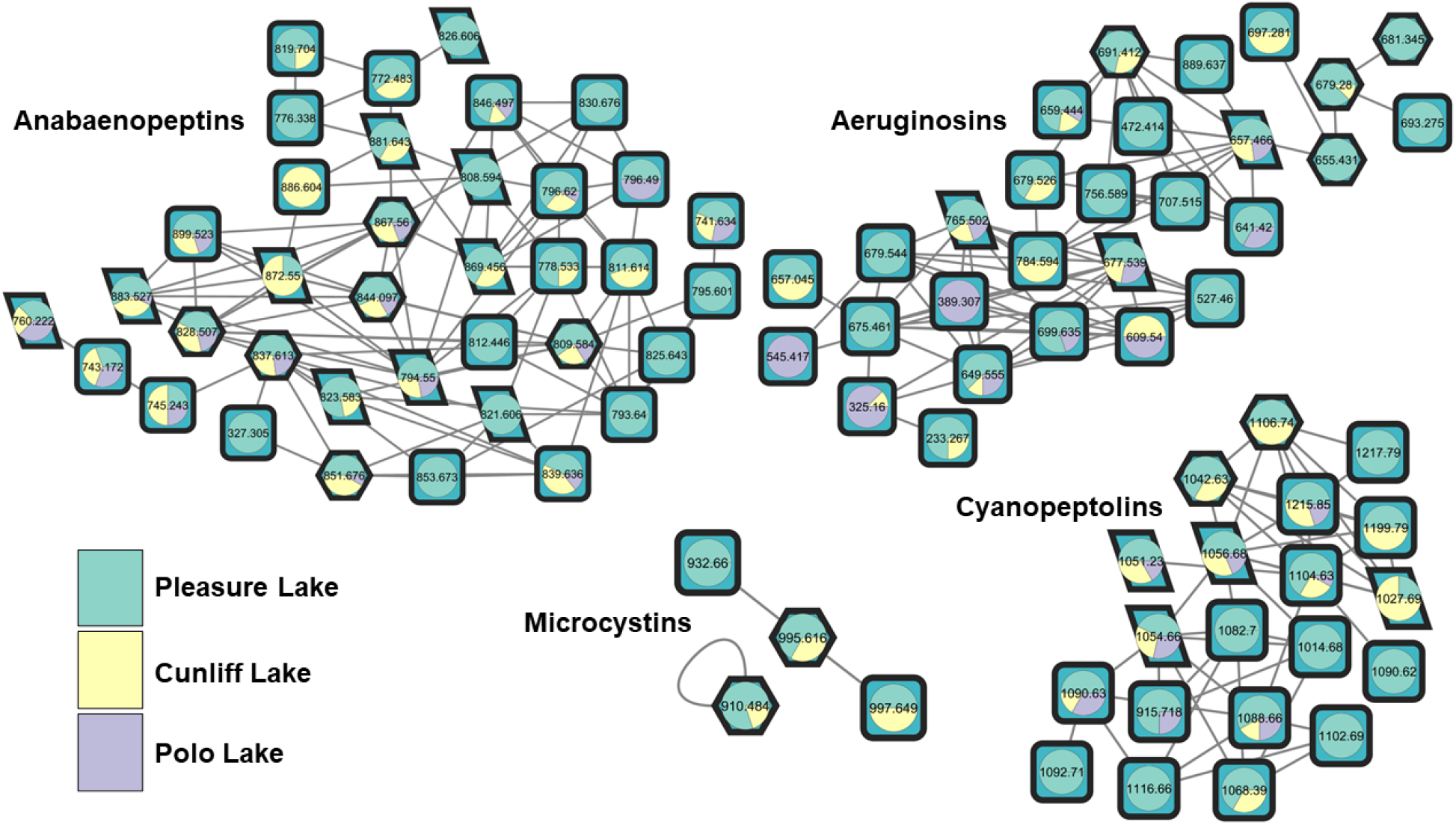
LC-MS/MS network of all extracts from surface water samples during the study period. Nodes are designated by their precursor *m/z* values and grouped into toxin classes. Pie slices indicate in which lake metabolites were detected (Pleasure, green; Cunliff, yellow; Polo, purple). Hexagons indicate that metabolites were amongst the 14 annotated in the study, while parallelograms indicate metabolites that are putatively known but were not specifically monitored. Squares indicate putatively new toxins.

**Figure 5.**
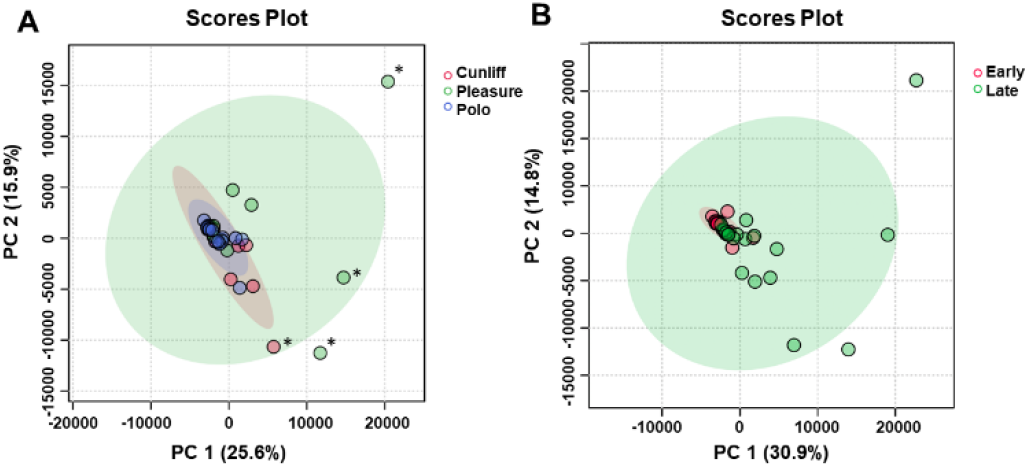
PCA analysis of bloom metabolites. (A) Comparison by site showed similarities in metabolite profiles among all sites. Asterisks show outliers (top to bottom: Pleasure 11/2/2022; Pleasure 11/9/2022; Cunliff 11/9/2022; Pleasure 11/30/2022. (B) Separating by early and late bloom periods showed differences in metabolite profiles.

Polo Lake was subject to phosphate control during the study period (Elizabeth Herron and Arthur Gold, personal communication). This treatment is designed to reduce free nutrients in the water column and trap them in the sediment. However, significant reduction in nutrients was not observed due to potential inputs from aeration or stormwater runoff. Thus, if nutrient control is ruled out, our current report provokes intriguing questions as to the ecological relationship between *Aphanizomenon* and *Microcystis* in terms of *Aphanizomenon*’s potential control of *Microcystis* growth and toxin production or if the relationship is primarily driven by temperature adaptation. While there are multiple studies pointing to allelopathic effects of *Microcystis* metabolites on *Aphanizomenon*,^11,47^ there is a gap in knowledge on the effect of *Aphanizomenon*’s metabolites on *Microcystis*. Future work will consist of co-culture experiments with these two organisms and the isolation, structure elucidation, and biological evaluation of putative new toxins from known classes and those from potentially novel classes. The potential aquatic toxicity and human health effects from emerging toxins and non microcystin toxins has been well described in fish models with non microcystin containing extracts and extracts from non-microcystin producing cyanobacterial strains still causing significant aquatic toxicity.^45,46^ Certain micropeptins have been shown to cause mortality and morbidity effects in fish models,^46^ and more research is needed to understand the full toxin suite in cyanoHAB events and their potential organismal effects.

The DNA metabarcoding approach showed interesting community structure changes over the course of the cyanoHABs. New hypotheses have formed – does *Aphanizomenon* produce allelopathic molecules to succeed *Microcystis* and other cyanobacterial species or alter the toxin production of other groups? Alternatively, are these community changes simply governed by temperature? Co-culture experiments could answer these questions. We identified several new toxins following molecular networking procedures and we have prioritized chromatography fractions for toxin isolation. Mixture analyses will be necessary to understand the combined effects of cyanotoxins on biological endpoints. Additionally, our data suggest that reducing cyanobacterial biomass in general rather than controlling for specific harmful groups will likely reduce all metabolites produced including toxins.

## ASSOCIATED CONTENT

### Supporting Information

The following files are available free of charge: Supporting Information with photographs and micrographs of cyanoHAB samples, chlorophyll a values, additional NMDS plots, and data on specialized metabolites correlated with cyanobacterial biomass (PDF).

## Author Contributions

M. B. and A. S. conceived the study. J. M., A. M. N. O., S. H., and H. X. performed data acquisition and conducted data analysis. B. J. provided materials and analysis and interpretation of data. M. B. wrote the manuscript with contributions of all authors. All authors have given approval to the final version of the manuscript.

## Funding Sources

We gratefully acknowledge NIH award 7R21ES033758-02 to M. B. and A. S.

## ACKNOWLEDGMENT

We are grateful to the members of the Stormwater Innovation Center for helpful conversations and about the lakes at Roger Williams Park. Acquisition of certain data in this publication was made possible by the use of equipment and services available through the RI-INBRE Centralized Research Core Facility at the University of Rhode Island, which is supported by the Institutional Development Award (IDeA) Network for Biomedical Research Excellence from the National Institute of General Medical Sciences of the National Institutes of Health under grant number P20GM103430.

## ABBREVIATIONS

ANOSIM: Analysis of Similarities
ASV: amplicon sequence variant
AUC: area under the curve
cyanoHAB: cyanobacterial harmful algal bloom
GNPS: Global Natural Products Social Molecular Networking
LC-MS: liquid chromatography-mass spectrometry
MC-LR: microcystin-LR
MS/MS: tandem mass spectrometry
NMDS: Non-metric MultiDimensional Scaling
PCA: principal component analysis
PETE: polyester track-etch

## Notes

https://massive.ucsd.edu/ProteoSAFe/dataset.jsp?accession=MSV000093676

### Competing Interest Statement

The authors have declared no competing interest.

