## Supporting Information for "Temporal Dynamics of Cyanobacterial Bloom Community Composition and Toxin Production from Urban Lakes"

**CONTENTS**

**Figure S1……………………………………………………………………………….3**

**Figure S2……………………………………………………………………………….4**

**Figure S3……………………………………………………………………………….5**

**Figure S4……………………………………………………………………………….6**

**Figure S5……………………………………………………………………………….7**

**Figure S6……………………………………………………………………………….8**

**Figure S7……………………………………………………………………………….9**

**Figure S8……………………………………………………………………………...10**

**Figure S9……………………………………………………………………………...11**

**Figure S10…………………………………………………………………………….12**

**Figure S11…………………………………………………………………………….13**

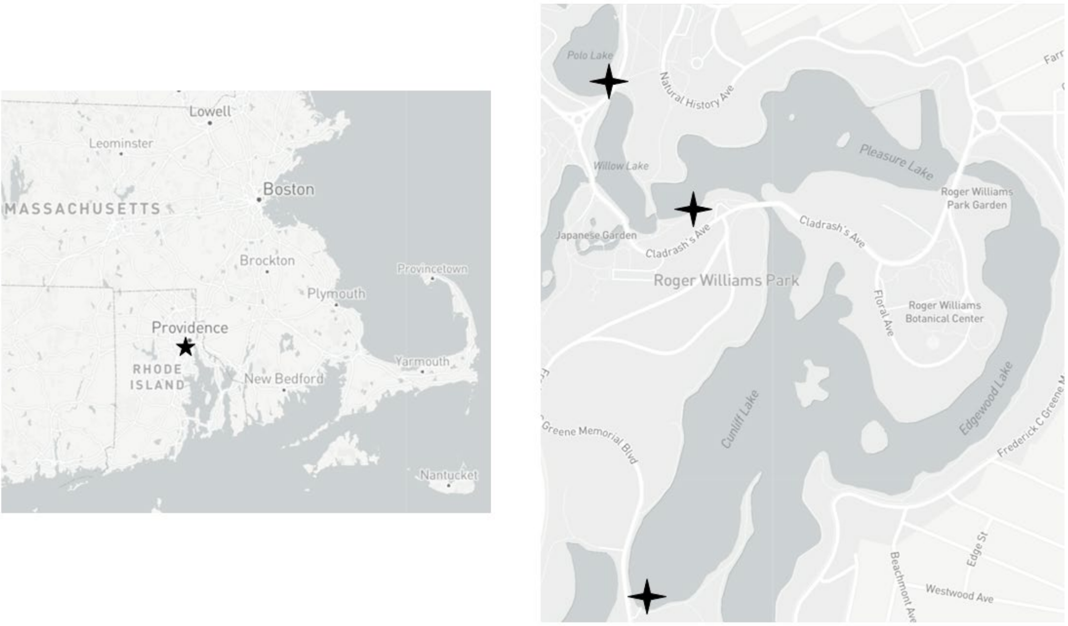


Figure S1. Map showing Providence, RI (left) and the three sampling sites at Polo Lake, Pleasure Lake, and Cunliff Lake (right).

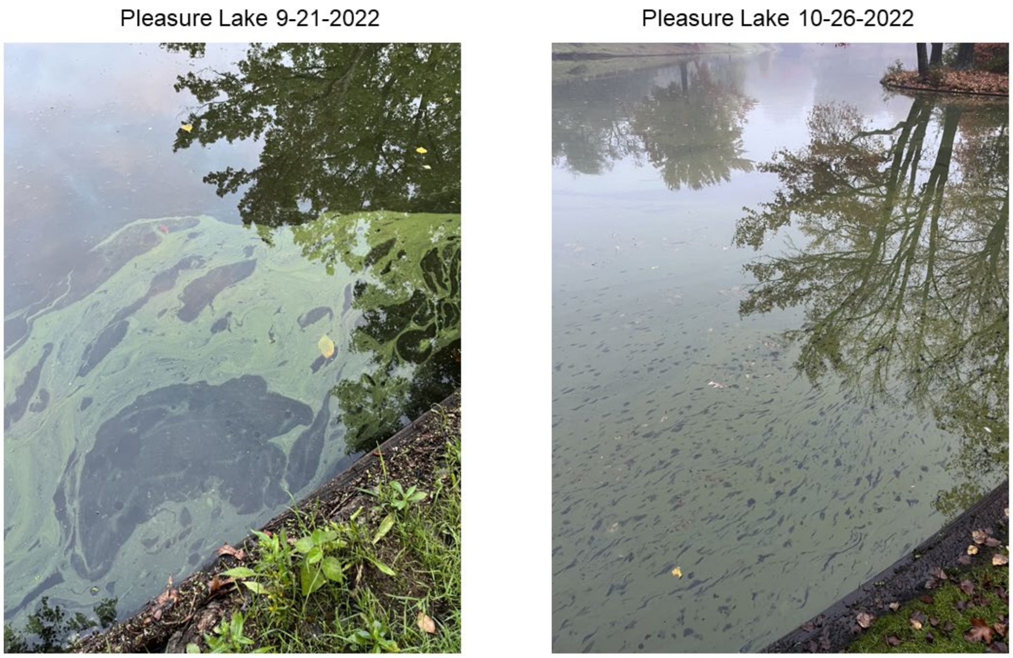


Figure S2. Photographs of surface water from Pleasure Lake on 9/21/2022 and 10/26/2022.

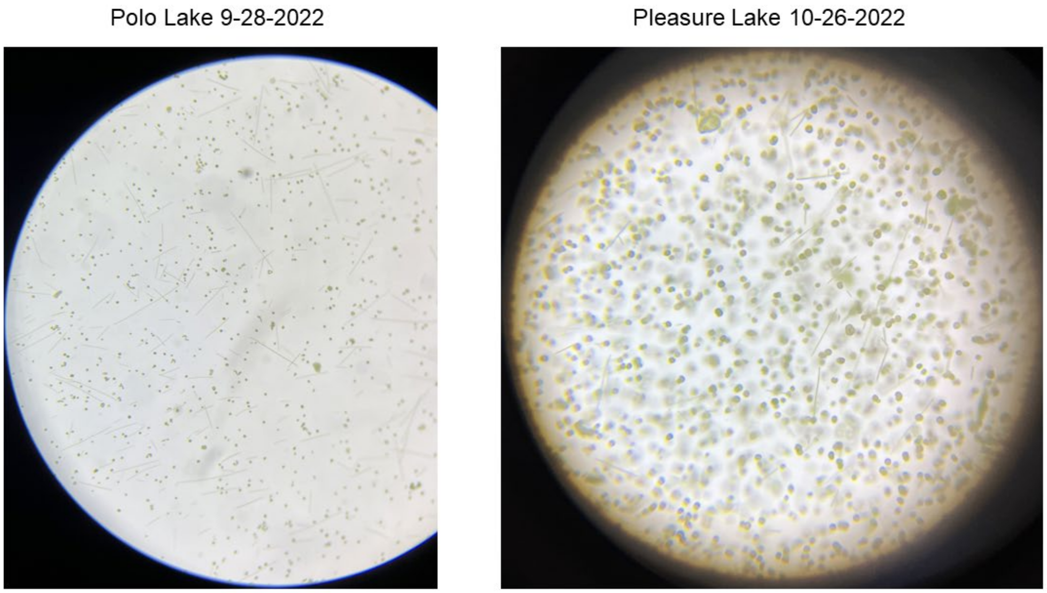


Figure S3. Dissecting microscope pictures of surface water samples from Polo Lake on 9/28/2022 and Pleasure Lake on 10/26/2022.

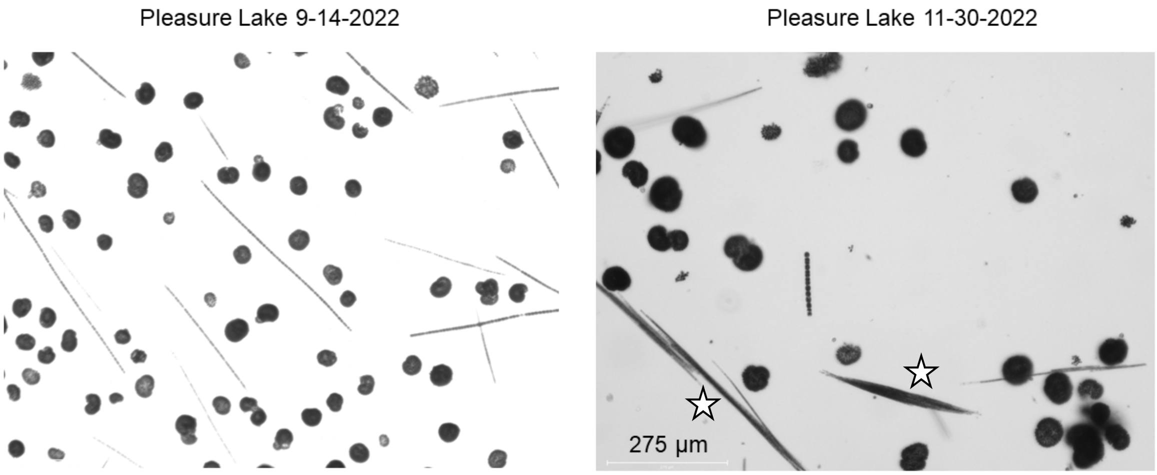


Figure S4. Photomicrographs of surface water samples from Pleasure Lake on 9/14/2022 and 11/30/2022. Stars show colonies of *Aphanizomenon* sp.

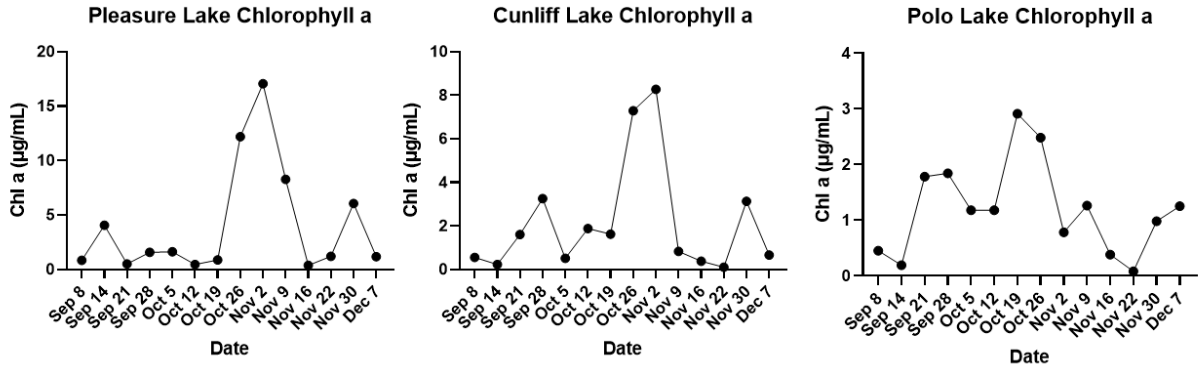


Figure S5. Chlorophyll a values (µg/mL) recorded from 9/8/2022 to 12/7/2022 at Pleasure, Cunliff, and Polo Lakes.

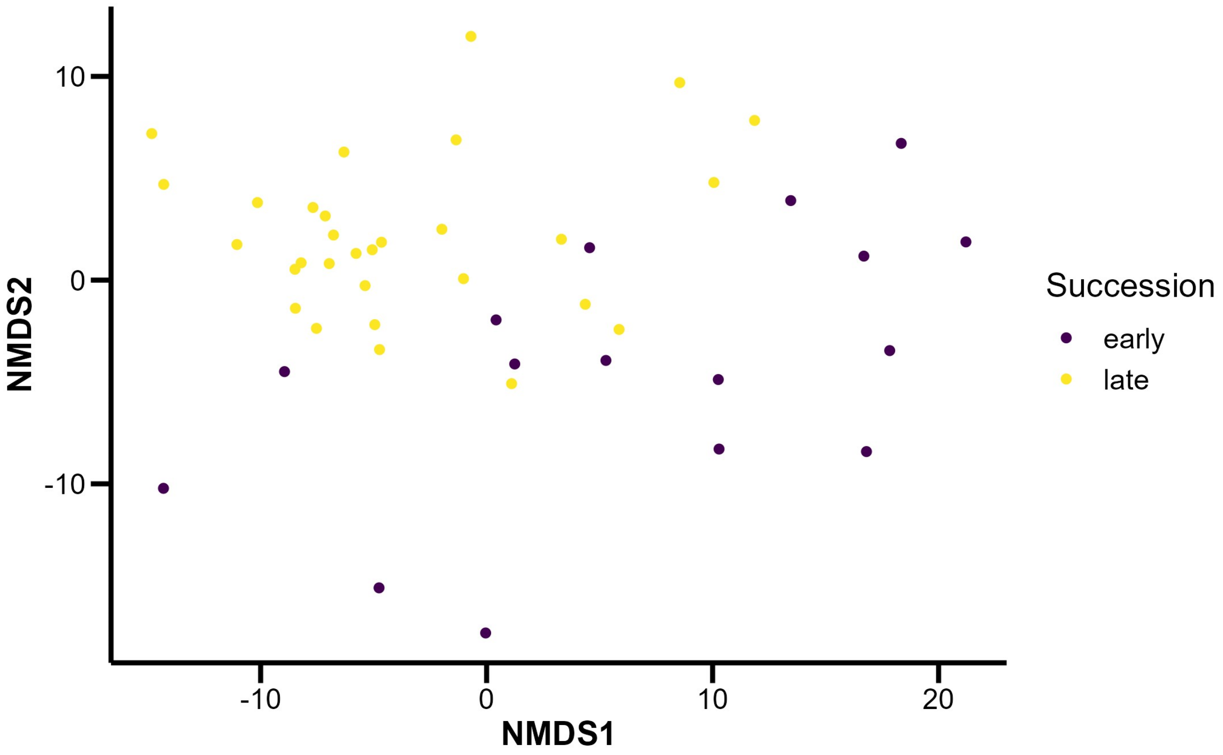


Figure S6. NMDS plots of the cyanobacterial taxa composition at all sites for early and late bloom periods. There was significant difference observed in the similarity of community composition in the two periods (ANOSIM, $p < 0.001$, R = 0.586).

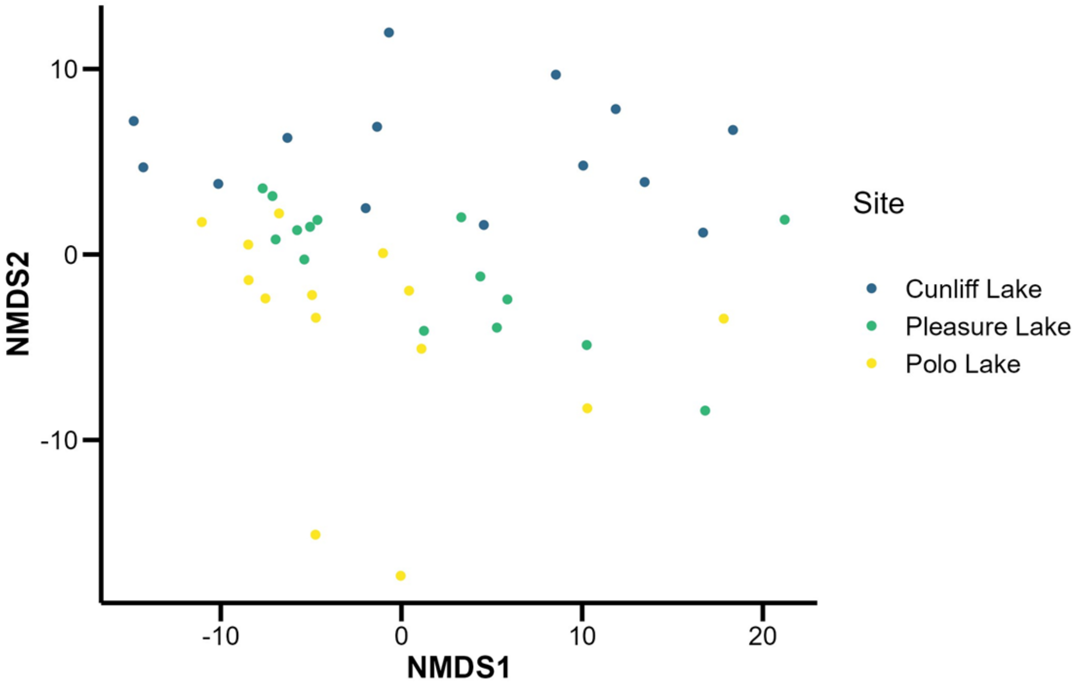


Figure S7. NMDS plots of the cyanobacterial taxa composition at all sites for the entire study period. There were significant differences observed in similarity of community composition amongst sites (ANOSIM, R = 0.14, $p < 0.01$).

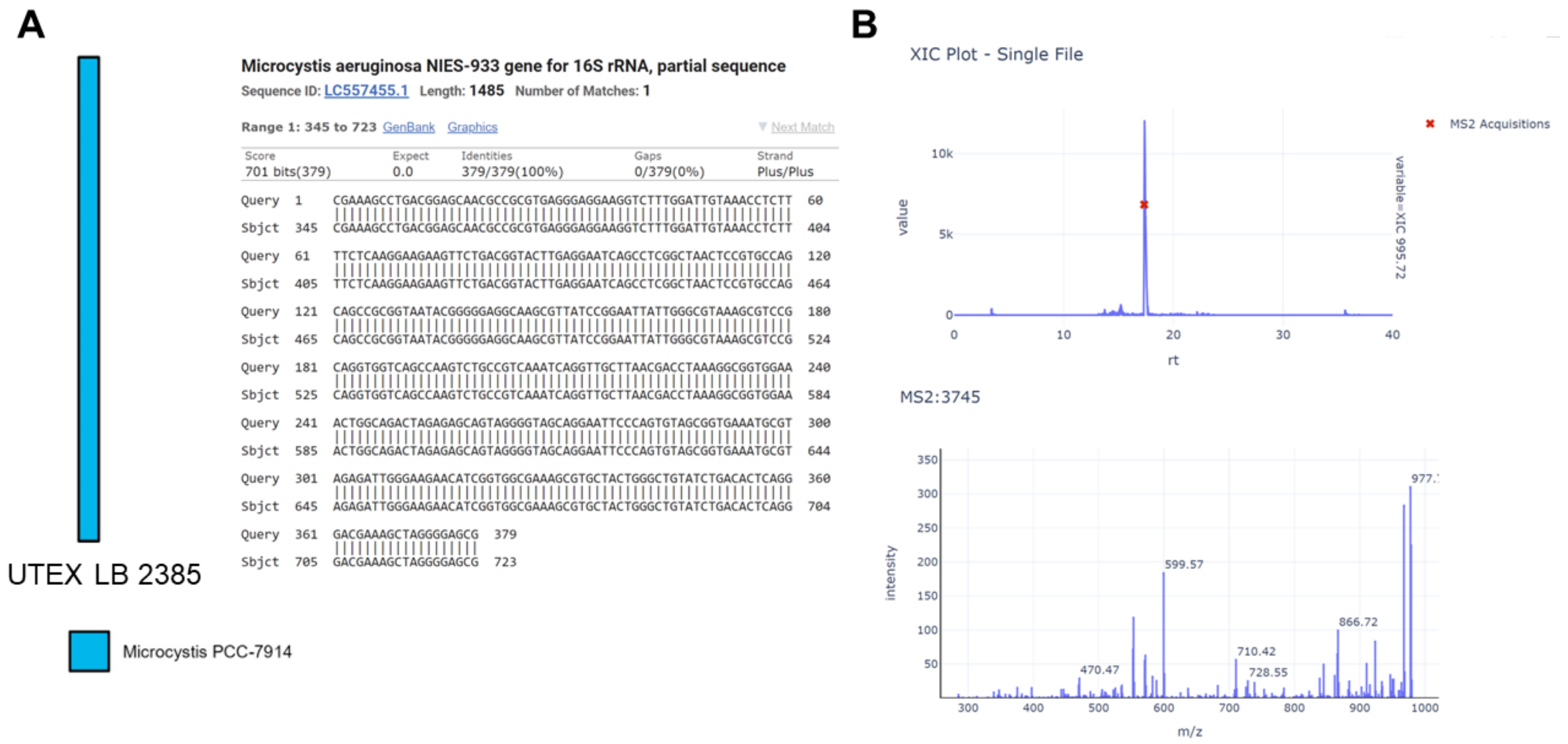


Figure S8. (A) ASV sequence analysis of the positive control *M. aeruginosa* culture showed that 100% of the ASVs were identified as *Microcystis*. BLAST searching showed that this ASV sequence showed 100% identity to that of *Microcystis aeruginosa* NIES-933, which is the strain (LB 2385) in the UTEX culture collection. (B) Extracts of positive control *M. aeruginosa* (UTEX LB 2385) showed that the strain produced MC-LR. An extracted ion chromatogram (XIC = 995.74) showed the precursor *m/z* for MC-LR (top panel) and the MS/MS fragmentation pattern matched that of MC-LR in the GNPS library (bottom panel).

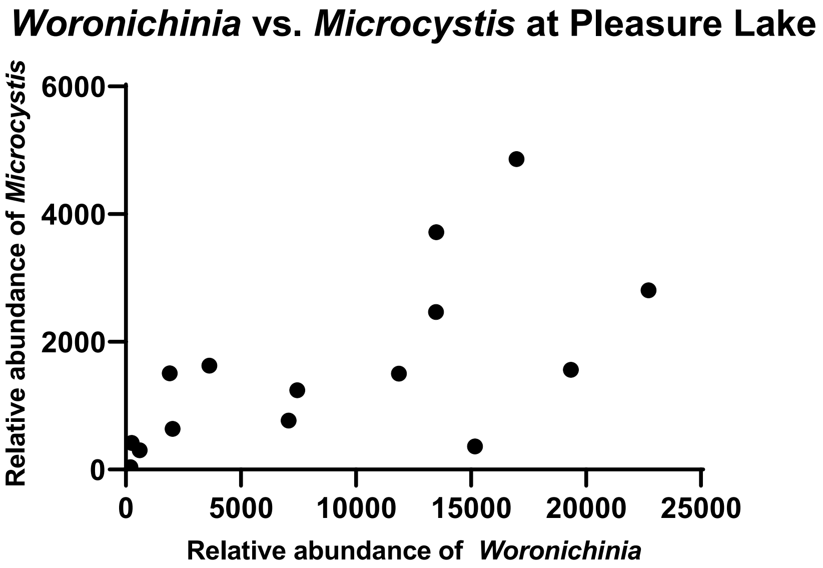


Figure S9. Correlation of *Woronichinia* and *Microcystis* relative abundance at Pleasure Lake measured via Pearson correlation coefficient ($p < 0.05$, $R^2 = 0.40$).

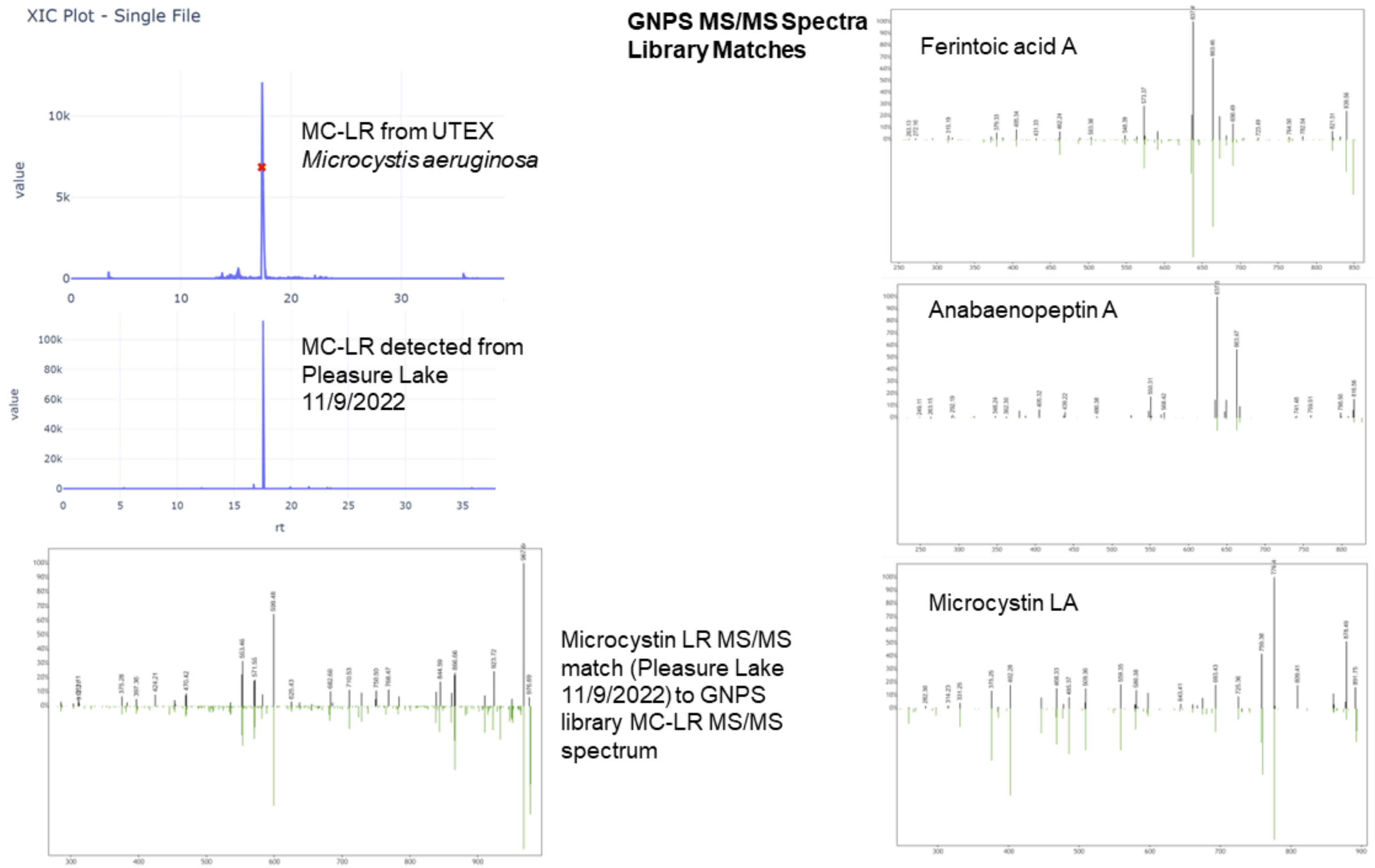


Figure S10. Annotation of cyanotoxins from environmental samples. Left panels show MC-LR retention time matches from two samples and MS/MS matches to the GNPS library. The rights panels show MS/MS matches to the GNPS library for ferintoic acid A, anabaenopeptin A, and microcystin LA.

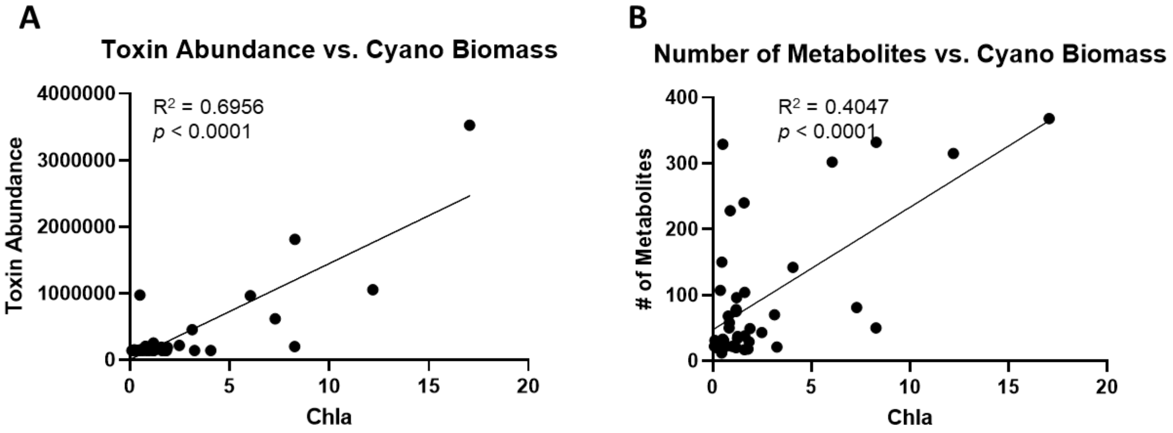


Figure S11. Toxin abundance (A) and the total metabolite number (B) in cyanobacterial bloom extracts significantly correlated with cyanobacterial biomass as measured by chlorophyll a values. Significance was determined using Pearson correlation coefficient in Prism (v. 9.5.1).
